# Effect of high-intensity anaerobic exercise on electrocortical activity in athletes and non-athletes

**DOI:** 10.1101/2024.08.29.610409

**Authors:** Élida Costa, Mariana Gongora, Juliana Bittencourt, Victor Marinho, Mauricio Cagy, Silmar Teixeira, Eduardo Nicoliche, Isabelle Fernandes, Danilo Fagundes, Caroline Machado, Juliana Dias, Renan Vicente, Pedro Ribeiro, Daya S. Gupta, Bruna Velasques, Henning Budde

## Abstract

**Aim:** The present study aims to verify the information processing in athletes through electroencephalography, analyze cortical areas responsible for cognitive functions related to attentional processing of visual stimuli, and investigate motor activity’s influence on cognitive aspects.

**Material and Methods:** The sample consisted of 29 subjects, divided into an experimental group (n = 13 modern pentathlon athletes) and a control group (n = 16 non-athletes). We collected the electrocortical activity before and after the Wingate Anaerobic Test. During the electrophysiological measures, the volunteers performed a saccadic eye movement paradigm. They also performed cognitive tasks, resting heart rate, and anthropometric measurements.

**Results:** A mixed ANOVA was applied to analyze the statistical differences between groups (athletes and control) and moments (before and after exercise) for F3, F4, P3, and P4 electrodes during rest one and task (pre-stimulus GO). There was an interaction for the group vs. moment factors in F3 [F = 17,129; p = 0,000; η² = 0.512], F4 [F = 22,774; p = 0,000; η² = 0.510], P3 [F = 11,429; p = 0,001; η² = 0.405], and P4 electrodes [F = 18,651; p = 0,000; η² = 0.379]. We found the main effect for group factors in the frontal and parietal electrodes of the right hemisphere (F4 and P4) and a main effect of the moment factor on the frontal (F3 and F4) and parietal (P3 and P4) electrodes. There was an interaction between the group vs. moment factors for the reaction time. The groups were different in Peak Power (Watts/kg), Average Power (Watts/kg), Fatigue Index (%), and Maximum Power (ms).

**Conclusions:** We identified chronic effects of exercise training on the cortical activity of modern pentathlon athletes, read-through differences in absolute alpha power, and acute effects of a high-intensity exercise session for athletes and non-athletes for electrocortical and behavioral responses.

## 1. Introduction

Sports can provide several motor and cognitive benefits. Athletes regularly practice sports activities professionally to improve performance, compete or maintain physical health, follow specific training routines, and participate in sports competitions (1–4). Studies with experienced athletes show the presence of different cortical activations that result in a “neural efficiency” brain function mechanism when compared with non-athletes (5,6). The “neural efficiency” hypothesis suggests that, as a result of sports training, expert athletes may exhibit cortical activation patterns seen through brain mapping techniques (7,8). Although several studies report the acute effects of exercise on cognitive functions (9,10). Few studies have investigated the underlying neural mechanisms in pentathlon athletes.

In a quantitative electroencephalography analysis (EEGq), there is an association of alpha rhythm with various states and functions, such as anticipatory attention, inhibitory control, information processing speed, memory, and sensory processing (11,12). Acutely, studies report changes in EEGq variables in the frequency domain. According to a literature review, absolute alpha power (AAP) increases after exercise compared to resting levels (13). Therefore, alpha rhythm may be a way to investigate cognitive functions.

Some studies have found relevant results about the effect of physical activity on attention to investigate cognitive processing. It is essential to clarify that physical activity is any motor action that takes the subject out of a state of suspension, and physical exercise is a structured and systematic activity that requires routine and planning (14). Llorens et al.(2015) reported that physical activity could modulate attention. Attention is a cognitive mechanism that allows someone to process information and relevant thoughts or actions, ignoring irrelevant or distracting stimuli (13,16,17). It can be subdivided into endogenous or voluntary attention and exogenous or reflex attention (18). Attentional and inhibitory control demands related to visuomotor responses are high in sports with open motor skills (19). In particular, saccadic eye movement tasks are relevant in measuring cognitive processes (20).

However, few studies have investigated the cortical changes produced by sports practice (17) and the acute effect of high-intensity exercise on the cortical responses of elite athletes, especially in sports practitioners with different visual-motor skills. Thus, an investigation into what occurs in cognitive processing in response to visual stimuli after a high-intensity exercise session may contribute to a better understanding of the influence of sports training on cognitive performance and the construction of training protocols.

The present study aims to verify the acute effects of exercise on information processing in elite athletes through electroencephalography. We investigated some cortical areas of sensorimotor integration (frontal and parietal cortex). Specific goals: 1) to investigate the acute effect of maximal anaerobic exercise in AAP; 2) analyze AAP at the frontal and parietal electrodes during the execution of a saccadic go-no-go task; 3) identify the differences in the AAP and the behavioral responses between athletes and non-athletes. Therefore, the hypotheses of the present study are: 1) modern pentathlon athletes would have a better response when performing a visual-motor task and higher AAP in sensorimotor integration areas when compared to non-athletes; 2) After a bout of maximal anaerobic exercise, both groups would have increased AAP in sensorimotor integration areas.

## 2. Material and methods

### 2.1. Participants

This experimental study comprised 29 volunteers divided into the experimental group (EG) and the control group (CG). The EC included 30 modern pentathlon athletes of the Brazilian Modern Pentathlon Team (4 females). The CG consisted of sixteen participants who did not engage in any sports activities (11 females) according to the Baecke questionnaire on physical activity (21). The volunteers were right-handed, regular, or corrected to normal vision and had no sensory, motor, cognitive, or attention deficits that could affect the saccadic eye movement task. Participants were not under the effect of any substance that could influence brain activity and had no history of psychiatric or neurologic disease. All participants answered an anamnesis, the Edinburgh Inventory (22), and the Structured Clinical Interview of the DSM-IV for Axis I Disorders (SCID-I) (23). They all provided written informed consent before the experimental procedures. This study was conducted following the guidelines in the declaration of Helsinki and was reviewed and approved by the Research Ethics Committee of the Federal University of Rio de Janeiro - UFRJ (protocol n°: 00996818.2.0000.5257) in February 2019. The authors did not have access to information that could identify individual participants during or after data collection. The final demographic characteristics and fitness data for the two groups are in Table 1.

**Table 1.**
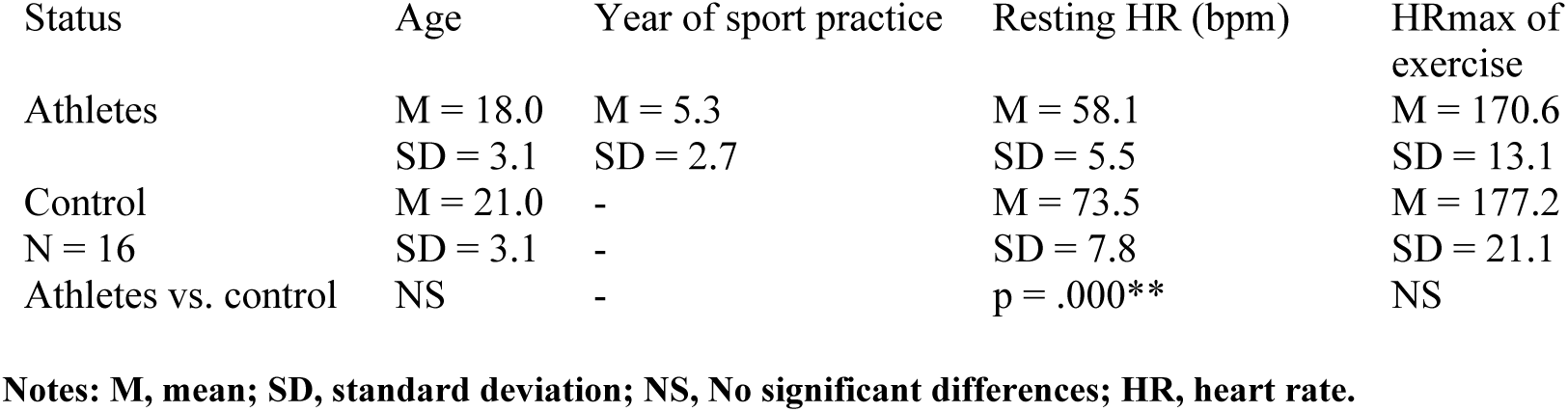
Participants’ characteristics.

### 2.2. Experimental procedure and task

The study design is represented in Figure 1 and consisted of the following stages: (i) psychometric; (ii) physiological assessments; (iii) experimental task before the anaerobic exercise (EEG 1); (iv) single session of anaerobic exercise; (v) experimental task after the anaerobic exercise (EEG 2) (Figure 1). The experiment occurred at the Biometrics Laboratory of the Physical Education and Sports School of the Rio de Janeiro Federal University (EEFD/UFRJ).

**Figure 1.**
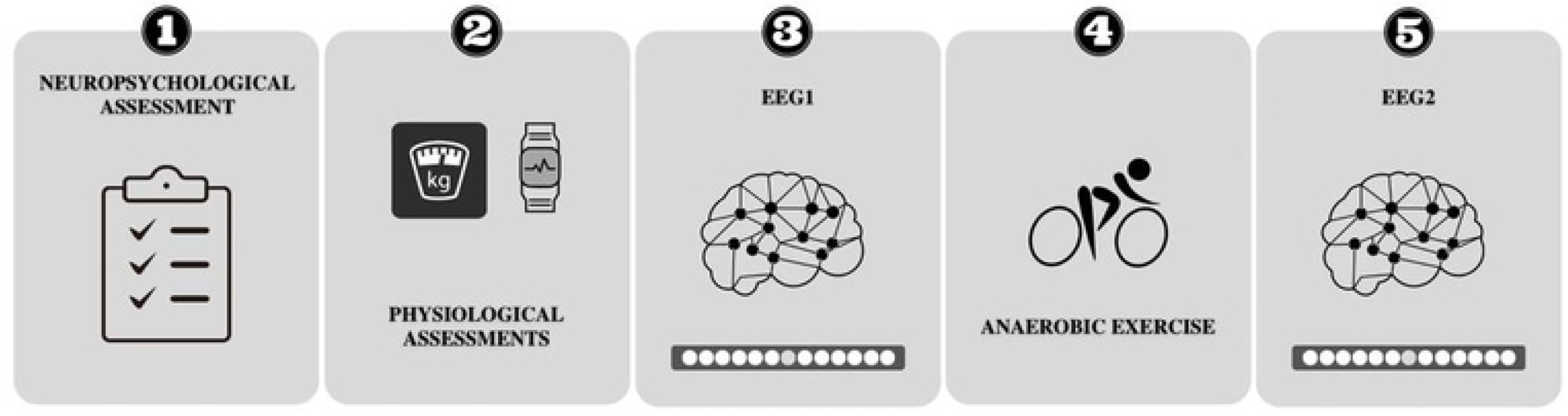
Experimental procedure. Participants performed the following stages: (1) neuropsychological and (2) physiological assessments. Each participant performed the electroencephalography acquisition during the saccadic eye movement task immediately before (3) and after (5) a single session of anaerobic exercise (4).

The neuropsychological assessments consisted of the following instruments: (i) Psychological Battery for Attention Assessment (PBA) (24); (ii) Five Digit Test (FDT) (25). The physiological assessments consisted of Heart Rate (HR) measurement monitored three times: at rest, during the anaerobic exercise session, and before EEG 2 (Polar® RS 800 CX - Polar Electro, Oy, Finland). Anthropometric weight and height measurements were collected to calculate the individual load for the exercise session (26).

Participants sat comfortably in a chair in front of a 120-centimeter bar of 13 light-emitting diodes (LEDs) positioned 100 cm above their eyes. Participants should follow the stimuli with their eyes, maintaining their heads still. Three colors on the central LED signaled each task phase: blue, green, and red. The blue LED was a warning and preceded the GO (green LED) or NO-GO (red LED). The green LED indicated that the participants should look for the target (LED-lit at one of the fixed points on the right or left). The red LED indicated that the participants should remain with their eyes fixed on the center of the bar. Each block consisted of 20 bands with a 50% distribution for the appearance of each stimulus (GO or NO-GO). Each block was composed of 20 attempts with a 50% distribution for the appearance of each stimulus (GO or NO-GO). The LED distribution was random (right or left) (Fig 2).

**Figure 2.**
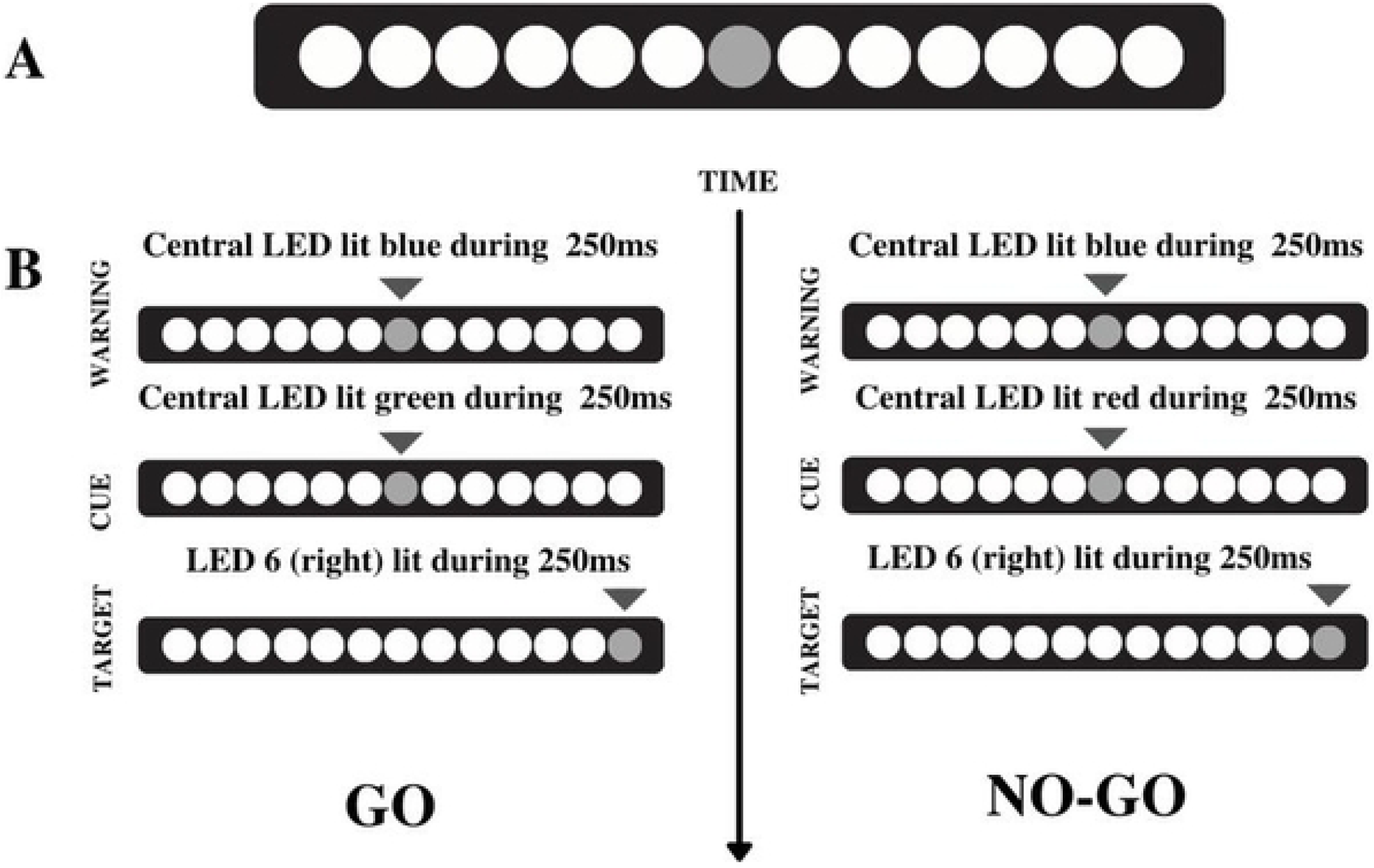
SEM task: this task starts with rest with eyes open for 3 minutes, five blocks with 20 stimuli (10 GO and 10 NO-GO) per block, and a final 3-minute rest. Each block was composed of 20 attempts with a 50% distribution for the appearance of each stimulus (GO or NO-GO). The distribution of the target LEDs was random (right or left).

### 2.3. EEG recording

The EEG signal acquisition was recorded using the 20-channel BNT-36 (EMSA – Medical Instruments, Brazil) EEG system and the SEM Acquisition. This program filtered the data: Notch (60 Hz), high-pass of 0.3 Hz, and low-pass of 80 Hz (order 2 Butterworth). Twenty electrodes were arranged on a lycra cap (EletroCap Inc., Fairfax, VA) along with the frontal, temporal, parietal, and occipital scalp regions, according to the 10/20 international system (27). Two more electrodes were positioned on the earlobes and set as a reference point, yielding 20 mono-pole derivations (using Fpz as a ground electrode). The caps were individually adjusted and put on each subject according to the participant’s head circumference and anatomical proportions. The signal corresponding to each EEG derivation came from the electric potential difference between each electrode and the pre-set reference (earlobes). The epochs were computed according to the stimulus appearance, four seconds before and four seconds after the stimuli. Each participant had ten epochs before the stimuli appearance and ten more epochs after the stimuli presentation.

The impedance levels for each electrode were kept below 10 kΩ. Two 9-mm electrodes in a bipolar montage estimated the ocular electric activity. The electrodes were positioned above and below the right eye orbit to register vertical ocular movements and on the external corner of the eyes to register horizontal ocular movements. We used the EEGLab program using Matlab 5.3® (The Mathworks, Inc.) to inspect the Visual artifacts.

### 2.4. Saccadic eye movement during EEG acquisition

Additional electrodes were positioned above and below the orbit of the right eye to record vertical eye movements (vEOG) and in the outer corner of the same eye to record horizontal eye movements (hEOG). The Neurophysiology and Neuropsychology of Attention Laboratory developed the saccadic software in DELPHI 5.0, which controlled the LED bar and determined the stimulus presentation.

### 2.5. EEG data processing

The offline preprocessing was performed using the EEGLAB software (http://sccn.ucsd.edu/eeglab) and included the following steps. Visual inspection and Independent Component Analysis (ICA) were applied to quantify reference-free data by removing possible sources of task-induced artifacts, such as eye blinking, temporomandibular muscle contraction, and body movements performed during data collection. Also, possible environmental noises may have occurred. ICA is an information maximization algorithm that derives spatial filters by blind source, separating the EEG signals into temporally independent and spatially fixed components (28). We excluded data from individual electrodes exhibiting loss of contact with the scalp or high impedances (> ten kΩ), as were data from single-trial epochs that exhibited excessive movement artifact (±100 µV). After the initial visual inspection, we applied the ICA to identify and remove any remaining artifacts, and we removed independent components resembling an eye blink or muscle artifact. The remaining components were projected back onto the scalp electrodes by multiplying the input data by the inverse matrix of the spatial filter coefficients derived from ICA. The ICA-filtered data were then re-inspected for residual artifacts. Epochs were selected between 1-sec pre-stimulus and 1.5-sec post-stimulus. The total number of epochs used after visual inspection and ICA for each group was as follows: non-athlete group (n 376); athlete group (n 353). We computed ERPs for the Fz, Cz, and Pz electrodes.

### 2.6. Anaerobic exercise

The Wingate Test of 30 seconds (26) was performed in an ergometric cycle (Monark® 828E, Stockholm, Sweden), imposed as a load of 0.075 kg · kg-1 of body weight and with an interface on the microcomputer. The ideal height of the bench was checked, adjusting to being close to 5° of knee flexion with the legs extended. During the test, all participants were verbally encouraged to use maximum strength in high intensity (> 85% of HRmax). The participants warmed up for 5 minutes at 60 bpm. Then, a 3-second sprint was performed before the load was activated to start the maximum anaerobic exercise for 30 seconds. Finally, the participants remained pedaling without load for 5 minutes for recovery, details in Fig 3.

**Figure 3.**
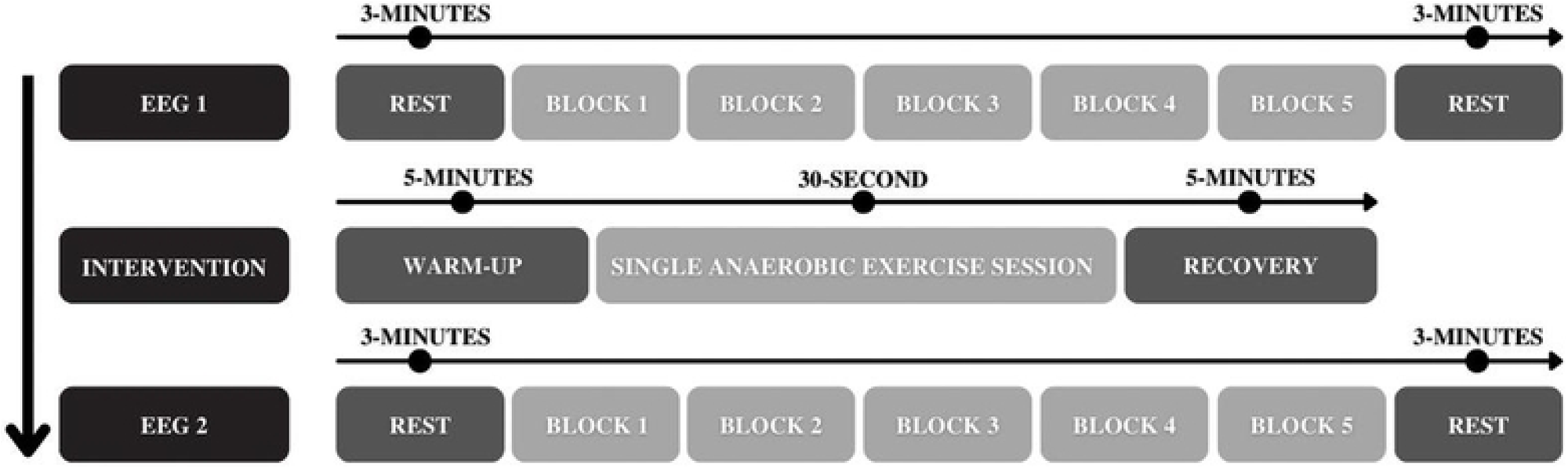
Details of the experimental design. EEG signal acquisition recorded during SEM before (EEG1) and after (EEG2) a single session of exercise: consisted of a previous task in rest with eyes open for 3 minutes, five blocks with 20 stimuli (10 GO and 10 NO-GO) per block, and final 3-minute rest. Intervention: a single session of anaerobic exercise (Wingate Test) consisted of 5 minutes of warm-up without load at 60 rotations per minute (rpm), followed by 30 seconds of pedaling at maximum intensity with load and 5 minutes of recovery pedaling without load.

### 2.7. Statistical Analysis

We used SPSS (SPSS Inc., Chicago, IL, USA), SigmaPlot (Systat Software Inc., Chicago, IL, USA), and Microsoft Excel for Windows (Microsoft, Redmond, WA, USA) to perform statistical analyses. We applied the paired Student’s t-test to determine the significance between groups for the following variables: concentrated attention, divided attention, alternate attention, inhibition, cognitive flexibility, resting HR, HRmax reached in the exercise test, HR before starting the EEG 2 and maximum power (ms) and fatigue index (Watts/kg). We used descriptive statistics with mean ± standard deviation (SD).

For electrophysiological measurements, our dependent variable was the absolute alpha power, and the independent variables were moment and group. A mixed ANOVA was applied to analyze the statistical differences between groups (athletes and control) and moments (before and after exercise) for F3, F4, P3, and P4 electrodes during rest one and task (pre-stimulus GO). We investigated the electrodes separately. The probability of 5% for type I error was adopted in all analyses (p<0.05).

## 3. Results

The present study examined absolute alpha power over the frontal (F3 e F4) and parietal (P3 e P4) electrodes. In the task condition, our results showed interaction for group vs. moment factors for F3 [F = 17,129; p = 0,000; η² = 0.512] (fig 4), F4 [F = 22,774; p = 0,000; η² = 0.510] (fig 4), P3 [F = 11,429; p = 0,001; η² = 0.405] (fig 5), and P4 electrodes [F = 18,651; p = 0,000; η² = 0.379] (fig 5).

**Figure 4:**
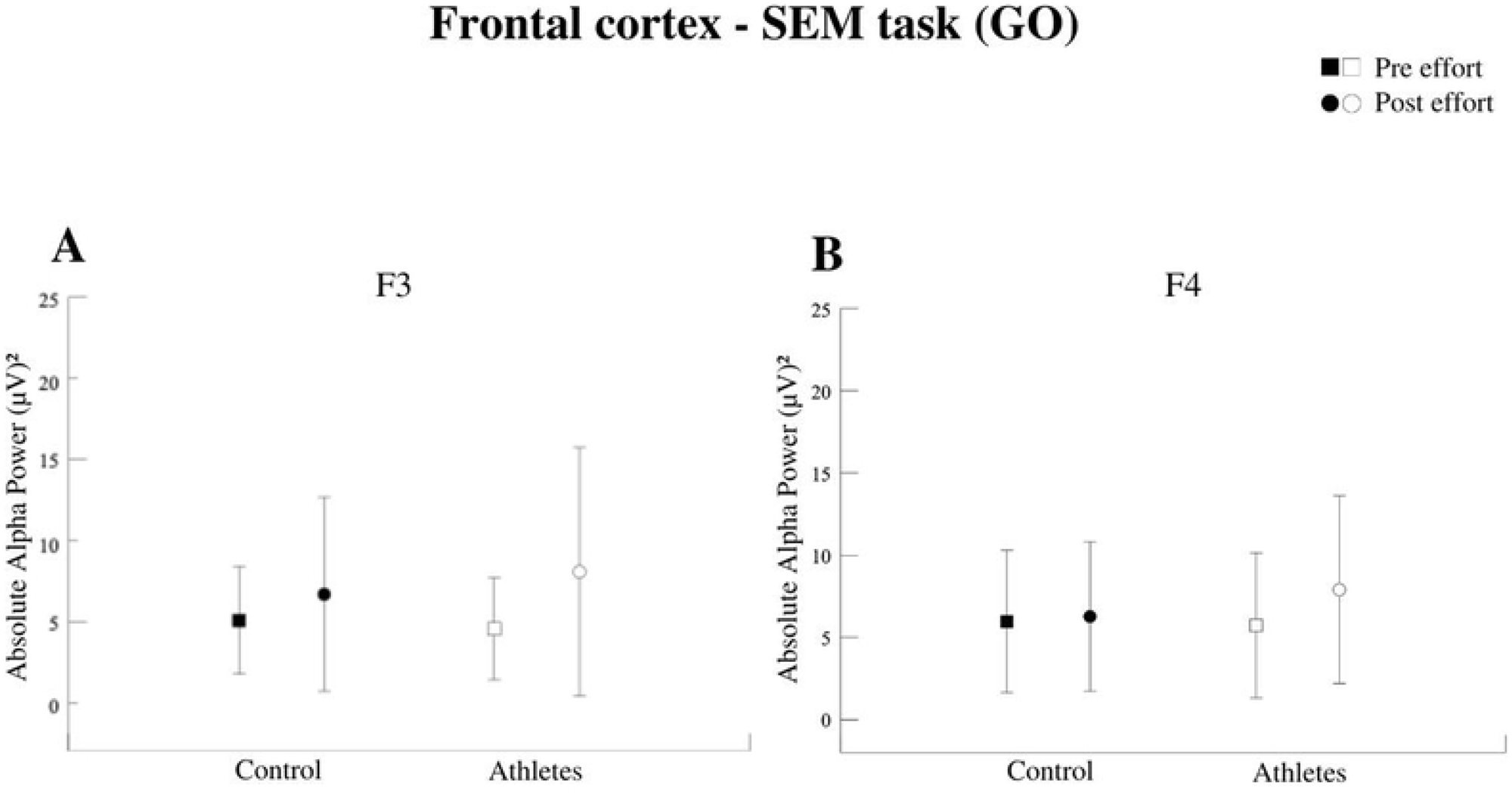
A- Absolute Alpha Power in the left frontal cortex (F3). Interaction between group vs moment factors (p = 0.000). Both groups increased the PAA after the effort. B - Absolute alpha power in the right frontal cortex (F4). Interaction between group vs moment factors (p = 0.000). The athletes had higher AAP than the pre-effort moment and greater AAP than the control after effort.

**Figure 5:**
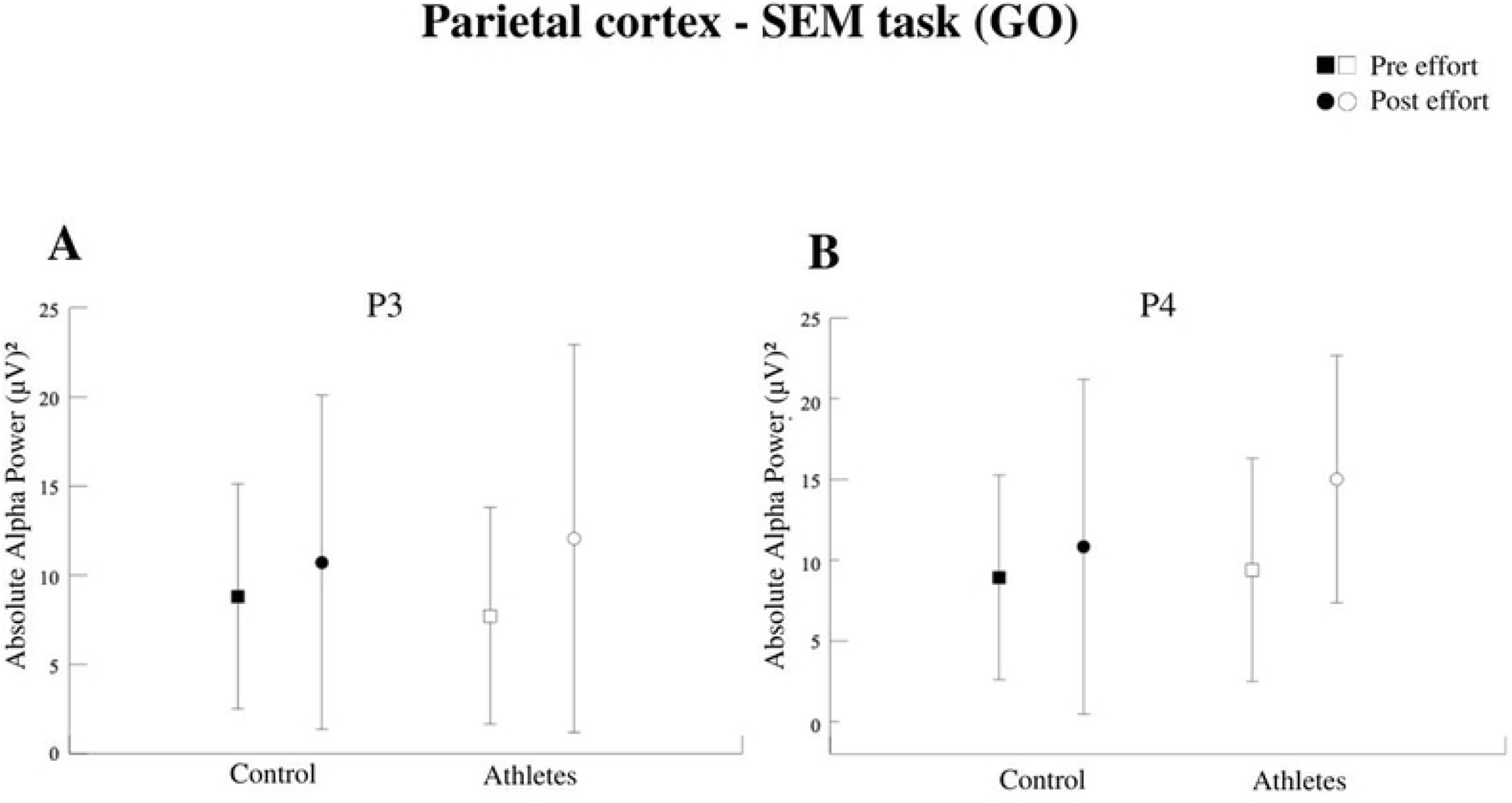
A – Absolute Alpha Power in the left parietal cortex (P3). Interaction between group and moment factors (p = 0.001). Both groups increased the AAP after the effort. B – Absolute Alpha Power in the right parietal cortex (P4). Interaction between group and moment factors (p = 0.000). Both groups increased the AAP after the effort. The athletes had a higher AAP compared to the control.

In the anaerobic exercise, statistically significant differences were found between the groups for Peak Power (Watts/kg) [F = 0.829; p = 0.019], (fig 7), Average Power (Watts/kg) [F = 0.312; p = 0.000] (fig 7), Fatigue index (%) [F = 0.091; p = 0.851] (figure 12) and Maximum Power (ms) [F = 0.245; p = 0.828] (fig 7). No significant differences were found between the groups for neuropsychological tests (table 2). In the SEM task, our results showed interaction for group vs moment factors for the reaction time variable [F = 8.291; p = 0.004], as well as the main effect for the moment [F = 28.082; p = 0.000], there was no SD for group [F = 0.003; p = 0.956] (fig 6).

**Figure 6:**
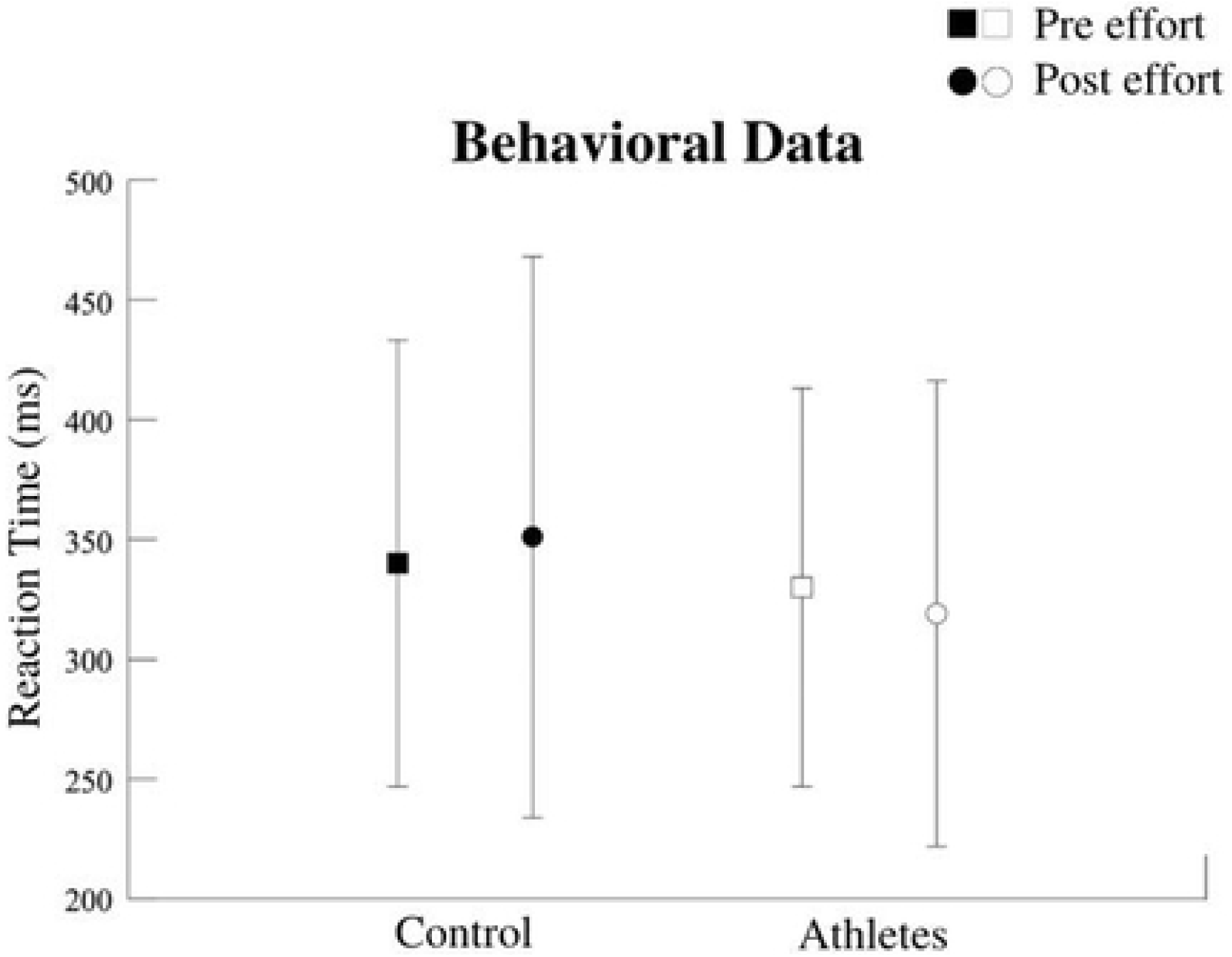
Reaction time (ms). Interaction for group vs Moment factors (p = 0.004). The athlete group reduced the RT after the intervention. The groups had no SD [F = 0.003; p = 0.956].

**Figure 7:**
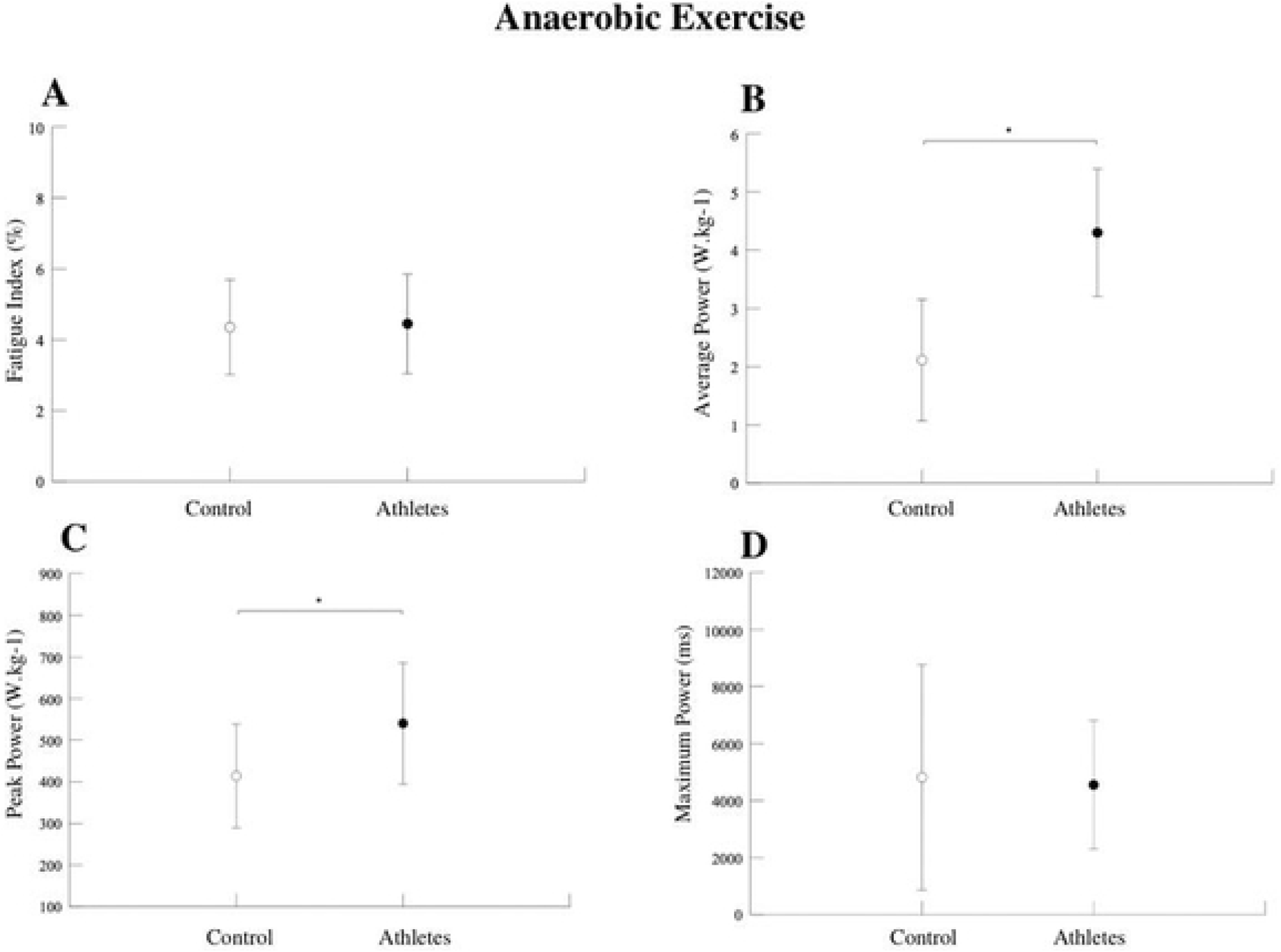
A - Fatigue Index Values (%). No significant differences were found between the groups (p = 0.851). B - Average Power Values (Watts/kg). Significant differences were found between the groups (p = 0.000). Athletes had higher Average Power compared to the control group. C - Peak energy values (Watts/kg). Significant differences were found between the groups (p = 0.019). Athletes had higher peak power compared to the control group. D - Maximum Power Values (ms). No significant differences were found between groups (p = 0.828).

**Table 2.** Neuropsychological dates.

| Status | Concentrated<br>Attention<br>(PBA) | Alternate<br>Attention (PBA) | Divided Attention<br>(PBA) | Cognitive<br>Flexibility<br>(FDT) | Inhibition<br>(FDT) |
| --- | --- | --- | --- | --- | --- |
| Athletes | M = 92.6 | M = 100.8 | M = 70.5 | M = 27.4 | M = 16.3 |
|  | SD = 19.5 | SD = 20.2 | SD = 23.2 | SD = 8.0 | SD = 6.3 |
| Control | M = 99.7 | M = 102.2 | M = 63.8 | M = 24.3 | M = 15.2 |
|  | SD = 15.3 | SD = 16.2 | SD = 23.4 | SD = 6.7 | SD = 5.6 |
| Athletes vs.<br>control | p = .265 | p = .826 | p = .428 | p = .259 | p = .622 |

Notes: M, mean; SD, standard deviation; PBA, Psychological Battery for Attention; FDT, Five Digit Test.

## 4. Discussion

Previous literature discussed physical exercise’s acute and chronic effects on cognition; however, few findings still report the neural mechanisms underlying behavioral responses. Thus, the present study aimed to verify physical exercise’s acute and chronic effects on information processing in modern pentathlon athletes through quantitative electroencephalography, analyzing absolute alpha power’s activity in sensorimotor integration cortical areas.

### 4.1. Acute effect of physical exercise

The effects of a single exercise session on cognitive processes and cortical responses may differ according to the intervention protocol or measures investigated. Previous literature investigated variations in the electrophysiological measurement of the alpha band (29). Alpha oscillations indicate a state of decreased cortical activation and are associated with relaxation and lower anxiety levels (30).

Niemann et al., (2013) Investigated the acute effects of a short session of intensive physical activity on cognitive performance in adolescents. The study found that a single intervention session could promote improved cognitive performance for individuals with higher levels of physical activity compared to individuals with lower levels. Furthermore, the results indicate that intensive physical activity had a beneficial effect on cognitive performance for all participants. Therefore, these data align with our hypothesis about the acute effects of physical exercise on cognitive performance. However, our cognitive test data did not show differences between the groups. Such findings may be related to the type of test used in our investigation.

In a literature review, Crabbe and Dishman (2004) demonstrated that the AAP increases throughout the scalp immediately after physical exercise. Therefore, such results corroborate our findings. Our results showed acute effects during the performance of the SEM task, with an increase in AAP for both groups in the left frontal hemisphere (F3) and parietal cortex (P3 and P4). AAP significantly increased after exertion for athletes in the right frontal hemisphere (F4). Schneider et al. (2009) demonstrated an increase in the alpha band for the Fp1 and F4 electrodes after a high-intensity exercise session in orienteering runners. These results corroborate our findings regarding the sample (athletes) and the increase in alpha power in the right frontal cortex (F4). Studies investigating the individual alpha peak frequency (iAPF) increased after intense exercise, and such results indicate a higher level of arousal and preparation for external stimuli (34,35). Therefore, alpha oscillations may be associated with better attention levels.

In a recent study comparing the effects of different exercise intensities on cortical activation in iAPF, acute physical exercise induces an increase in iAPF, a cortical parameter associated with the speed of neural information processing. However, when observing the responses at different intensities, the results revealed a significant increase in iAPF after high-intensity exercise (85-90% HR max). In addition, subsequent measurements showed that such changes persisted for about 30 minutes after an exercise interruption (35). These findings reinforce our results regarding alpha frequency band oscillations as an acute effect of high-intensity exercise. Thus, it seems possible to conclude that the findings converge to changes in the alpha frequency band caused by a single exercise session, regardless of the intensity or measure of the investigated alpha frequency band.

Our results also demonstrated an acute effect for improving the information processing speed measured through the visuomotor reaction time. The group of athletes reduced the RT after the effort, consistent with the previous results that demonstrated an impairment of RT responses during high-intensity exercises, with a supposed slowness in processing information. However, McMorris et al. (2011) reported specific results for different exercise intensities measured after exercise. They showed improved cognitive performance after moderate-intensity exercise and impairments after high-intensity exercise, measured through reaction time and accuracy. These studies compare the effects of different exercise intensities on the RT responses. Budde et al. (2010) found a significant improvement in adolescents’ working memory after a low-performance exercise session (50–65% of HRmax) (37). When discussing the effects of different exercise intensities, it can be seen that light exercise plays a role in hippocampal memory function (38). However, some terms, such as “very light intensity,” may not be used accurately in the studies (39). Thus, higher intensities and repetitions have been recommended to promote beneficial effects on cognitive functions (39,40). However, there is a need to understand exercise-based cognitive enhancement strategies better and the specific dose-response relationship and investigate the duration of exercise-induced effects (41). Our results do not cover these variables, so we do not measure other possible responses. In addition, the heterogeneity of the experimental methods used in the reviewed studies still needs help understanding the topic.

One of the possible explanations for the divergences in the literature on behavioral and neural results after an acute exercise session may be the time between the end of the exercise and the acquisition of the second investigation measure. Therefore, we performed our measurements after the return of HR to previously established parameters (110% of HRrep) and not immediately after effort, and some of the studies pointed out in the discussion investigated the results immediately after effort (42) or during exercise (43,44). Lambourne and Tomporowski’s (2010) results are consistent with our results measured after an exercise session on the cycle ergometer. Budde et al., (2008) demonstrated benefits in concentration and attention of adolescents after an intervention with coordinative exercises. The study concluded that this type of exercise can induce a pre-activation of cortical areas responsible for attentional functions. Therefore, the variation in results indicates the complex dose-response relationship between exercise and cognition.

Our results demonstrated that the increase in AAP is an acute effect of exercise, suggesting that AAP can be a biomarker of better performance in attention tasks; once the alpha band increases in sensorimotor areas are associated with a higher level of excitation and preparation for external stimuli (46). In addition, the electrophysiological results demonstrated a reduction in the visuomotor reaction time, which suggests an improvement in the speed of processing the stimulus.

### 4.1. Chronic effects of physical exercise

Previous studies have shown different cortical activity patterns among athlete subjects compared to beginners or non-athletes. EEGq measures these changes and is a possible marker of specialization for athletes, being part of the theoretical bases that reinforce the efficiency of the neural hypothesis (47–50). Ludyga et al., (2019) Investigated the effects of aerobic and coordination training on children’s behavioral and neurophysiological measures of inhibitory control through 10-week intervention protocols. No differences were found regarding the improvement of inhibitory control. Therefore, the practice time seems fundamental to observing neurophysiological changes.

Koutsandréou et al., (2019) comparing different types of exercise, it was observed that preadolescent children’s working memory performance benefited from both cardiovascular and motor exercise programs. However, the increase in working memory performance was more significant for children in motor exercise than cardiovascular or control exercise. These data highlight another variable that impacts the observed cognitive responses. Another essential variable to consider is age. Studies suggest that levels of executive function improve during adulthood and, therefore, different mechanisms impact performance in early and late life, which may cause changes in electrophysiological markers and performance in attention and inhibitory control tasks (53,54).

We found significant differences between the groups of athletes and non-athletes, especially for the task analyses. A significant increase in the AAP was identified for the group of athletes concerning the control after the stress test in all areas of the right hemisphere (F4 and P4). Consistent with our findings, Duru and Assem (2018) reported that when evaluating the realization of a cognitive workload paradigm, karate athletes demonstrated a higher alpha band power in the parietal and occipital cortices than non-athletes.

Experiments comparing professional versus novice dancers showed that professional dancers had more alpha synchronization in the mid-parietal, parietotemporal, and parietooccipital brain regions than novice dancers while performing an imagination paradigm (56). Therefore, previous studies are consistent with our findings on increasing AAP for experienced athletes performing a task with cognitive demand.

Alpha oscillations in the parietal region are lower in anxiety than in customary conditions, suggesting an inverse relationship between the alpha frequency band and anxiety levels (57). Furthermore, the increase in alpha power in areas of the parietal cortex has been interpreted as better levels of focused attention and savings in energy expenditure for information processing, being associated with higher performance indices in specialist athletes, demonstrating that training is capable of improving problem-solving (47). Thus, it is possible to conclude that our results are consistent with our study and reinforce the neural efficiency hypothesis for elite athletes, which is when skilled individuals exhibit a relatively different brain activity response called neural efficiency. Studies with elite athletes compared to non-athletes demonstrate different neural responses when subjected to the same experimental task. Such results are read in previous literature as adaptive and efficient mechanisms (5,6).

Among the limitations that make it difficult to discuss and interpret data consistently is the heterogeneity of the studies reported in the literature since the studies present divergent methodological procedures, including sample, experimental design, and investigated electrophysiological measurements. Budde et al. (2012) examined the effect of an intermittent maximal exercise intervention on selective and sustained attention tasks in a group of physically active adults. The results indicated that more physically active participants performed better on cognitive tests after the intervention. Thus, higher physical activity levels may lead to underlying neurobiological adaptations that regulate cognitive processes. Exercise variables such as exercise prescription according to external load, internal load, and influencing factors may interfere with interindividual response variability in neurocognitive measures (59). Thus, Herold et al., (2021) suggest that future studies investigating aspects of cognition and physical exercise should be rigorous in their experimental designs. More studies are needed, with samples from different sports, control of other variables, and even a longitudinal protocol to more efficiently study the chronic factors of sports training.

The high AAP indices follow the neural efficiency hypothesis since their interpretation provides information on cortical energy expenditure. Thus, there is an inverse relationship between the increase in the alpha band in some cortical regions and the activity of groups of neurons (60), which is interpreted as cortical energy saving for information processing.

## 5. Conclusion

Modern pentathlon athletes had greater absolute alpha potency than non-athletes, which suggests the chronic effects of sports training on cortical activity. Regarding the acute effects of a high-intensity exercise on electrocortical activity, an increase in absolute alpha power was found for both groups and a decrease in reaction time for athletes only.

Therefore, electrocortical data indicated that experienced athletes have a different pattern of cortical activity than non-athletes, which can be read as less activation in the information processing areas (frontal and parietal cortices), which aligns with the neural efficiency hypothesis. Furthermore, our results suggest that a single physical exercise session can modify neural responses that converge to improve cognitive performance.

Such findings contribute to justifying the practice of sports before cognitively demanding tasks. This study has limitations that need to be considered when interpreting the findings. It was not possible to expand the sample to other groups. The study’s cross-sectional nature is one of the main limitations, as the lack of sample control does not allow us to measure the effects of sports training on the cortical functions of athletes. Longitudinal studies are needed to explore the role of sports training on electrocortical responses, including different groups. Moreover, exercises at different intensities can be designed to investigate acute neural responses. Thus, we suggest that future studies examine the effects of different exercise intensities in other groups.

